# Evaluating the capabilities and challenges of layer-fMRI VASO at 3T

**DOI:** 10.1101/2022.07.26.501554

**Authors:** Laurentius (Renzo) Huber, Lisa Kronbichler, Rüdiger Stirnberg, Philipp Ehses, Tony Stöcker, Sara Fernández-Cabello, Benedikt A. Poser, Martin Kronbichler

## Abstract

Sub-millimeter functional imaging has the potential to capture cortical layer-specific functional information flow within and across brain systems. Recent sequence advancements of fMRI signal readout and contrast generations resulted in wide adaptation of layer-fMRI protocols across the global ultra-high-field (UHF) neuroimaging community. However, most layer-fMRI applications are confined to one of ≈100 privileged UHF imaging centers, and sequence contrasts with unwanted sensitivity to large draining veins. In this work, we propose the application of vein-signal free vascular space occupancy (VASO) layerfMRI sequences at widely accessible 3T scanners. Specifically, we implement, characterize, and apply a cerebral blood volume (CBV)-sensitive VASO fMRI at a 3T scanner setup, as it is typically used in the majority of cognitive neuroscience and clinical neuroscience fMRI studies. We find that the longer 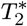, and stronger relative *T*_1_ contrast at 3T can account for some of the lower z-magnetization in the inversion-recovery VASO sequence compared to 7T and 9.4T. In the main series of experiments (N=16), we test the utility of this setup for motor tasks and find that-while being limited by thermal noise-3T layer-fMRI VASO is feasible within conventional scan durations. In a series of auxiliary studies, we furthermore explore the generalizability of the developed layer-fMRI protocols for a larger range of study designs including: visual stimulation, whole brain movie watching paradigms, and cognitive tasks with weaker effect sizes. We hope that the developed imaging protocols will help to increase accessibility of vein-signal free layer-fMRI imaging tools to a wider community of neuroimaging centers.

**Graphical abstract:** 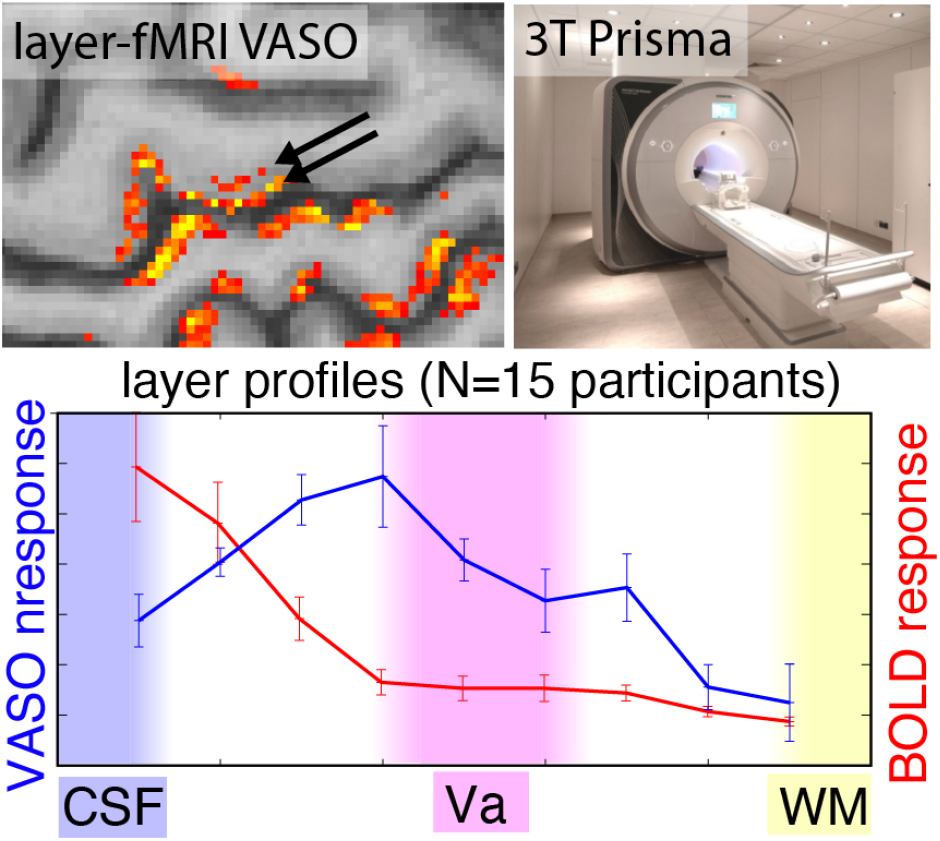

## 1. Introduction

Recent methodological advances in fMRI contrast and readout strategies allow researchers to approach the mesoscopic regime of cortical layers. This enables a mapping of cortical information processing within and across brain systems.

Layer-fMRI has the potential to provide circuitry information of neural processing that was previously only accessible with invasive procedures in animal research (Yang et al. 2021). Until now however, most layer-fMRI studies have been confined to specialized ultra-high field (UHF) equipment (7-9.4 Tesla) and venous-biased sequences, which is problematic (Bollmann and Barth 2020; Weldon and Olman 2020). This limits layer-fMRI to fulfill its potential in becoming a useful research tool for cognitive and clinical neuroscience. Specifically:

1. The need for UHF scanners limits layer-fMRI applications to ≈100 MRI labs globally and prohibits the widespread adoption of layer-fMRI as a neuroscientific imaging tool. Layer-fMRI at 3T (Akin and Özen 2019; Kim and Ress 2017; Knudsen et al. 2022; Koopmans et al. 2010; Lifshits et al. 2018; Markuerkiaga et al. 2020; Olman et al. 2007; Puckett et al. 2016; Ress et al. 2007; Scheeringa et al. 2016; Taso et al. 2021; Wu et al. 2018) can increase the availability of layer-fMRI worldwide by two orders of magnitude. However, only about 8% of all the human layer-fMRI papers used 3T (source: https://layerfmri.com/papers/).
2. The conventional fMRI contrast of gradient-echo (GE) BOLD imposes unwanted signal bias in the superficial layers (Olman et al. 2007). This complicates the neural interpretability of fMRI signal changes in the superficial layers. The additional application of cerebral blood volume (CBV)-sensitive vascular space occupancy (VASO) methods can mitigate such biases (Hua et al. 2013; Lu et al. 2003). VASO is a non-invasive fMRI sequence approach that is sensitive to changes in CBV changes by means of an inversion-recovery contrast generation. In VASO fMRI, an inversion-recovery pulse sequence is used to selectively null out blood water magnetization at the image of the image acquisition, while leaving extra-vascular signals for detection. An increase in CBV during task-evoked neural activation is then associated with an overall MR-signal decrease, which in-turn is believed to be proportional to the volume increase of nulled blood.

In this study, we aim to address both constraints with a new imaging methodology of a 3T-optimized VASO layer-fMRI sequence that utilizes a 3D-EPI readout (Poser et al. 2010; Stirnberg and Stöcker 2021).

While layer-fMRI VASO has been successfully applied in dozens of 7T research labs, its limited detection sensitivity has so far discouraged researchers to test its feasibility at 3T. Here, we want to argue that, theoretically, it should be quite straightforward to translate layer-fMRI VASO from 7T to 3T. Which is not the case for conventional BOLD:

- GE-BOLD is based on a susceptibility contrast that linearly scales with the field strengths (Martindale et al. 2008). Additionally at 7T, BOLD benefits from the generally increasing SNR increase with multi-channel coils at higher field strength of a factor 3.3 between 3T and 7T (Pohmann et al. 2016). This means that (in the thermal noise dominated regime of layer-fMRI) the voxel-wise BOLD detection sensitivity is reduced by a factor of 7.7, going from 7T to 3T. This is a large SNR penalty and can be a limiting constraint for BOLD at 3T. VASO, on the other hand, is a *T*_1_-based contrast. And thus, VASO does not suffer from the reduction of the susceptibility contrast at lower field strengths. Hence in VASO, the expected sensitivity reduction is ‘only’ in the order of a factor 3.3.
- Fig. 1A) depicts that 3T VASO can benefit from the stronger blood-tissue *T*_1_-contrast compared to 7T. While blood and tissue *T*_1_-values decrease with weaker field strengths, the relative difference between blood and tissue increases. Depending on the repetition times and flip angles used, this can boost the available relative GM magnetization at the blood-nulling time, which ultimately increases the VASO detection sensitivity at 3T compared to 7T. This effect can account for the lower absolute z-magnetization at 3T compared to 7T (see Fig, 1A).
- Layer-fMRI VASO at 7T is challenged by the fast 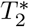 signal decay during relatively long EPI readouts with large imaging matrix sizes. At 3T, the 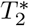 decay is slower, which helps maintaining the detection sensitivity across a longer echo train length,=. Further more, the slower decay also helps to maintain the sharpness of the imaging point spread function (see Fig. 1B). Depending on the echo time and readout duration, this can boost the VASO sensitivity at 3T compared to 7T.
- High-resolution EPI comes along with large readout trains and, thus, it can be limited by a low bandwidths (in the phase encoding direction). This makes layerfMRI at 7T prone to phase-inconsistency artifacts and to intermittent ghosting (example in Fig. 1C). At 3T, such artifacts are mitigated, which ultimately increases the layer-fMRI sensitivity at 3T compared to 7T.

**Fig. 1).**
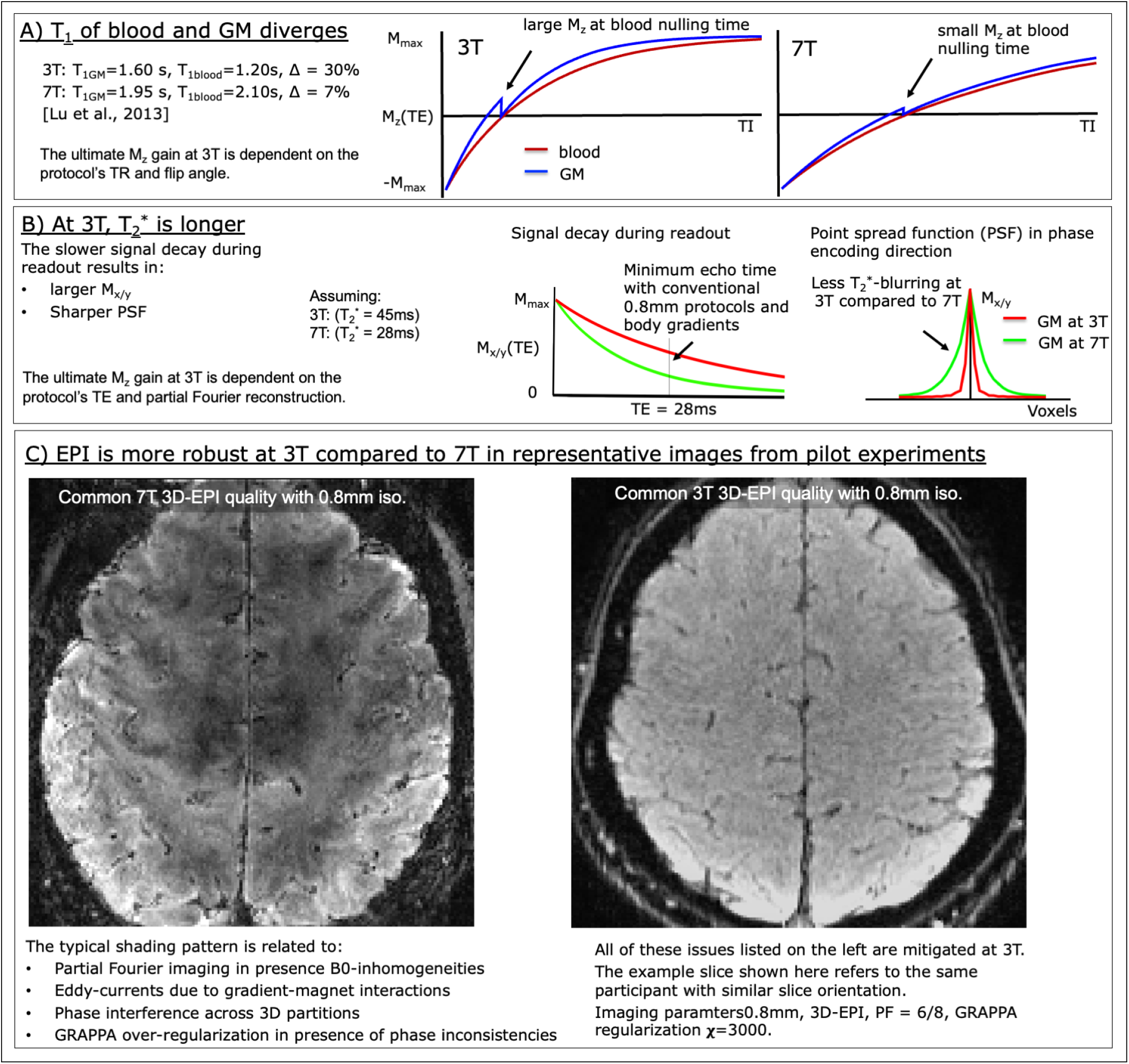
expected advantages of 3T over 7T for layer-fMRI VASO. Panel A). At 3T, the relative difference between GM and blood *T*_1_ relaxation is stronger than at 7T. This is despite the fact that *T*_1_ values increase with field strengths. The stronger *T*_1_ contrast at 3T results in larger *M*_z_-magnetization (in units of *M*_0_) at the blood nulling time. Panel B). At 3T, the slower 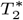-decay results in reduced signal decay during the readout. With the relatively large matrix-sizes in layer-fMRI (>200), this can boost the transverse magnetization *M_x/y_* (in units of *M*_0_), and result in sharper images. Panel C). Typical EPI image quality at 7T and 3T. The *B*_0_, and *B*_1_ inhomogeneity at 7T results in phase errors and signal brightness variations. These artifacts can be temporally insatiable and limit the fMRI detection sensitivity. Such artifacts are reduced at 3T.

The goal of this study is to implement, characterize, and validate an imaging methodology of 3T CBV-weighted layerfMRI. We seek to quantify the feasibility of layer-fMRI VASO at 3T for the test case of motor tasks (main study). And furthermore, we aim to perform exploratory supplementary studies to investigate the generalizability of the developed protocols for a larger range of study designs including: visual stimulation, whole brain movie watching paradigms, and cognitive tasks with weaker effect sizes. Lastly, we aim to compare the protocol across 3T, 7T and 9.4T and investigate potential 3T-specific intra-vascular BOLD contaminations.

## 2. Methods

N=16 participants were scanned on a SIEMENS Prisma (64ch coil) with a 3D-EPI sequence developed by Stirnberg and Stöcker (2021) for VASO imaging with whole-brain MAGEC capabilities (Huber et al. 2021a). All procedures carried out in this study were approved by the Ethics Review Committee for Psychology and Neuroscience (ERCPN) at Maastricht University (ERCPN-180_03_06_2017) and the Ethics Committee of the Paris-Lodron-University of Salzburg (EK-GZ 20/2014), following the principles expressed in the Declaration of Helsinki.

Across all 3T experiments, the participants were placed in the scanner bore, such that the imaging region is close to the isocenter. The RF transmit birdcage body coil of the Prisma scanners used here has a length of 50cm. The VASO inversion pulse was played out ‘globally’ (without a slab-selective gradient). This means that the inversion slab thickness was determined by the size of the transmit coil. And, given the head positioning at the iso-center, the inversion was effectively confined to the participant’s head and neck regions. In the SS-SI VASO approach used here, an inversion slab of at least 14cm is necessary to mitigate inflow effects of fresh blood (Huber et al. 2014b). However, the inversion slab should be small enough to allow refilling the ‘fresh’ blood between inversion pulses (here 4.7s). Thus, it was relevant in here that the blood in the heart was outside the inversion slab. We used an adiabatic Hypersecant 6 pulse for inversion.

### 2.1. Main scanning protocol (motor)

A slab protocol (26 slices) was used for motor experiments. Imaging parameters were: FOV 177mm, isotropic nominal resolution of 0.82mm, TE=28.5ms, *TR_shot_*=80.6ms, transmit reference voltage=250V, read bandwidth = 772 Hz/Px, echo spacing=1.4ms, partial Fourier=6/8, *k_y_* GRAPPA 3 (effective echo spacing=0.47ms, phase encode bandwidth=9.9Hz/Px). An inversion delay of 550ms was used to acquire the k-space center of the VASO imaging readout approximately at the *M_z_*-nulling time of once-inverted (nonsteady-state) blood. Variable flip angles were used to mitigate *T*_1_-relaxation related blurring in the partition direction: flip angles = 33.1°-60.0°. Water-selective excitation was done with binomial 1-1 pulses having a bandwidth time product of 8. One initial external phase navigator scan was acquired and applied globally (Stirnberg and Stöcker 2021). The acquisition of each complete k-space volume was *TR_vol_*=2.075s. This means that it took *TR_pair_*=4.7s to acquire a pair of BOLD and VASO images. All images were reconstructed using a generic vendor-preinstalled GRAPPA (Griswold et al. 2002) implementation compatible with 2D CAIPIRINHA (Breuer et al. 2006) using (‘IcePAT’) (Kellman and McVeigh 2005) with a 3D GRAPPA kernel of *k_x_*=5, *k_y_*=4, *k_z_*=3. The slab orientation was axial and covered the superior part of the primary motor cortex BA4B (see Fig. 2A). Complex-valued signals of all 64 RF receive channels were combined with the vendor provided software Prescan Normalize method (Jellúš and Kannengießer 2014), which can be advantageous in areas close to small coil elements (Schmitt and Rieger 2021). We used the specific adaptive combine algorithm EVD_PSNPC (eigenvalue decomposition with adjust coil sensitivity phase correction). This protocol was tested on 16 participants.

**Fig. 2).**
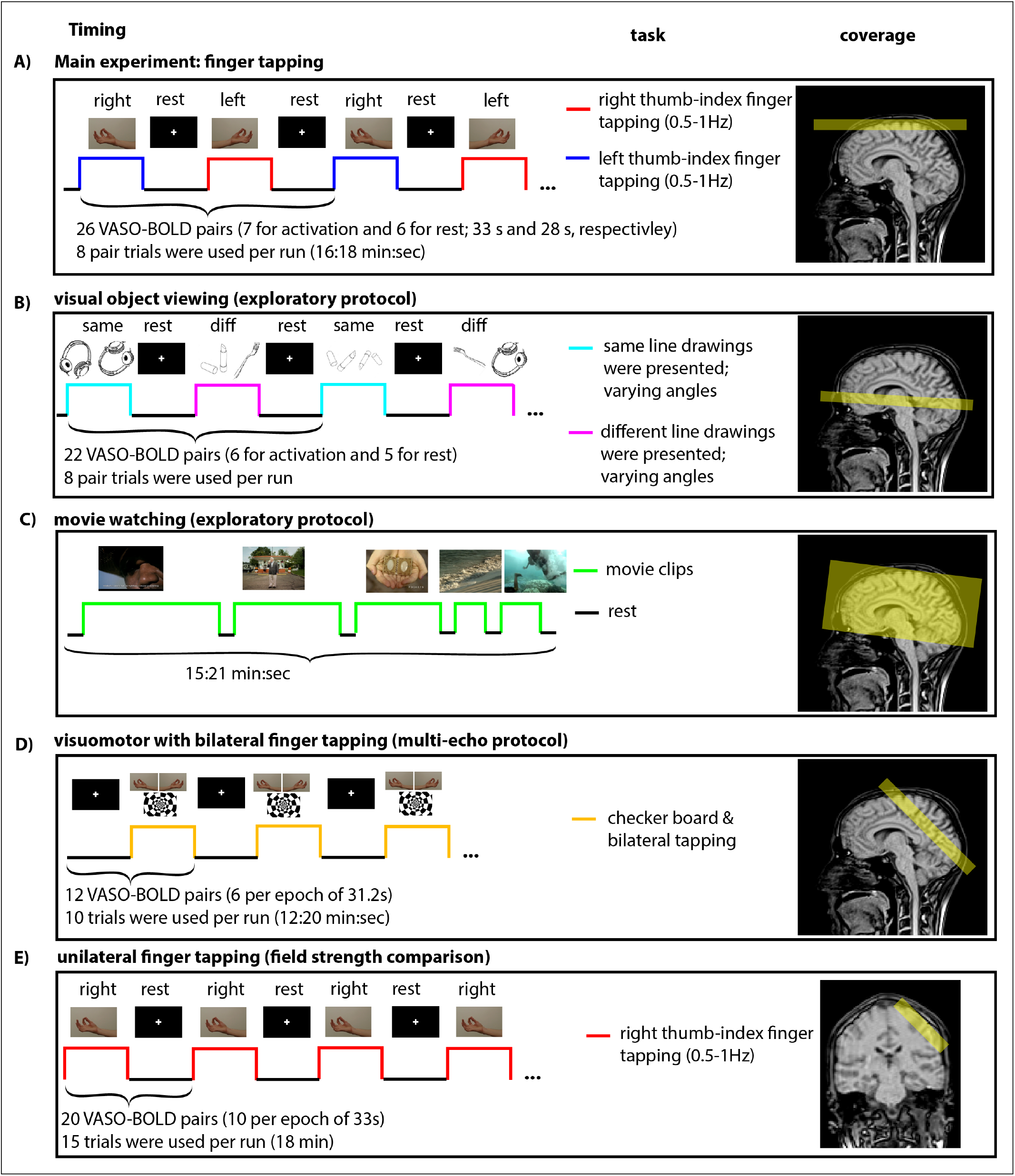
task timings and coverages used for the main experiments and supplementary experiments.

### 2.2. Other scanning protocols for exploratory purposes

In order to explore the generalizability of the results from motor tasks (described above), we extended it for other types of experiments. This includes layer-fMRI in visual areas, whole brain protocols, multi-echo protocols, and one field strength comparison.

#### 2.2.1. Visual protocol

For visual stimulation experiments, we applied almost the same slab imaging parameter as described above. The only major difference was that the slab position was lowered to cover the calcarine sulcus (Fig 2B). Further minor differences were: number of slices was 28, which increased the *TR_pair_* to 5.1s. The slab orientation was axial-like with a bit of tilting. Data were acquired in 16 participants, however, only one representative dataset was analyzed and is shown here.

#### 2.2.2. Whole brain protocol

In order to explore the feasibility of whole-brain layer-specific connectivity analyses, we extended the slab protocol described above to 120 slices for an imaging coverage large enough to capture the entire cortex in conventional head sizes. Since the readout of so many slices would be longer than the *T*_1_-recovery, we spread out the acquisition of a complete set of k-space partitions across four inversion-recovery periods with effective inversion times of 1.2s and 2.5s, for VASO and BOLD, respectively (acquired every 3.5s). We used an in-plane segmentation factor of 2 with an effective shot-specific readout duration of *TR_shot_* = 43ms (effective echo spacing of 0.23ms, phase encode band-width of 19.8Hz/Px). The echo time was 15.6ms. In addition to the in-plane acceleration of 3, we applied acceleration of 2 in the second phase encoding direction (total undersampling factor of 6). This protocol was tested in two sessions.

#### 2.2.3. Multi-echo protocol to assess BOLD contaminations

To assess the effectiveness of BOLD-correction in VASO sequences, additional two participants were invited and scanned with segmented multi-echo protocols. Protocol parameters were: skipped-CAIPI 6 · 1 × 3_*z*2_ sampling without z-blips (Stirnberg and Stöcker 2021) (no *k_y_*-GRAPPA, *k_z_*-GRAPPA 3, 6 in-plane segments with *k_z_* CAIPI shift 2), one inversion per 3 segments, two echo times (12ms, 48ms), 0.9mm isotropic. Image matrix size 212×216×12 with a FOV of 200mmx200mmx11mm. Four volumes were acquired every 5.2s (effective temporal resolution): VASO_*TE*1_, VASO_*TE*2_, BOLD_*TE*_1, BOLD_*TE*2_. 6 runs (12 min each) were obtained im two participants.

#### 2.2.4. Field strengths comparison in one participant

While the resolution and the task trial timing in the main experiment is designed to provide results that can be compared to previous layer-fMRI VASO studies at 7T and 9.4T (Huber et al. 2018), these experiments have been conducted with different participants. In order to have a more direct comparison across field strengths, we reached out to previous participants that had already conducted layer-fMRI VASO experiments at 9.4T and 7T. One participant volunteered to repeat the same experiment at 3T with the newly developed layer-fMRI protocol. In accordance with previous 7T and 9.4T experiments, the imaging field of view was solely covering the motor cortex of one hemisphere with a tilted imaging slab aligned perpendicular to the central sulcus (see Fig. 2E).

### 2.3. Stimulation and Task

#### 2.3.1. Main experiment with finger tapping task

The motor task consisted of 16 min alternating activation-rest blocks. Periods of activation and rest were alternated every 14 and 12 volume acquisitions for activation and rest, respectively (32.9s, 28.2s respectively). During activation periods, participants were instructed to perform a thumb-index finger pinch movement every 1-2 seconds. Each 16 min run was executed twice per session.

#### 2.3.2. Supplementary tasks

To test the generalisability of the motor results for the visual cortex, we also used a visual task. Namely, we used a visual repetition priming design, in which line drawings of objects were presented. In repetition epochs, always the same object picture was presented. Whereas in alternating epochs, different object pictures were used. In both types of epochs, the visual angle of subsequent visual stimuli was alternated to reduce low-level visual cortex adaption.

To test the generalisability of the motor results for whole brain protocols, we also used movie watching tasks. Namely, we used the 15 min collection of movie clips that are established for exploring advanced fMRI methodology from the 7T HCP study (aka MOVIE1). This collection consists of 5 separate short stories (1:03-4:05min) interspaced with 20s of rest. The clips are from independent films freely available under a Creative Commons license and are entitled: Two Men (2009), Welcome to 221 Bridgeville (2011), Pockets (2008), Inside the Human Body (2011), 23 Degrees South (2011) and LXIV (2011).

To characterize the echo-time dependence of the BOLD-corrected VASO signal, we used a visuomotor task, consisting of a full-field flickering checkerboard. The participants were instructed to perform a finger tapping task as fast as they can with both hands, whenever they see a flickering checkerboard. Activation and rest periods were alternated every 12 TRs (2.6s each).

For comparisons across field strengths, the trial timing of the tapping task from the main experiment described above was modified to match with the timing of previous experiments (Huber et al. 2018). Namely, we used a unilateral tapping of the right hand only, with activation-rest periods of 33 sec, each. This was repeated 15 times.

### 2.4. Offline data processing

#### 2.4.1. Data conversion

Dicom images from the scanner were sorted according to the ICE-dimension of ‘sets’ and converted to nii time series with isisconv (Enrico Raimer, Der Orfa, MPI Leipzig) using the script here: https://github.com/layerfMRI/repository/blob/master/conv/conv_Kronbichler.sh.

#### 2.4.2. Motion correction and BOLD correction

Concomitantly acquired time series consisting of blood-nulled and BOLD contrasts were separately corrected for motion using SPM12 (Functional Imaging Laboratory, University College London, UK). Motion-corrected time series were corrected for BOLD contaminations, by means of dynamically dividing blood-nulled signals with not-nulled BOLD signals. In order to mitigate non-steady state effects, the division was performed on a two-fold temporally upsampled time series. This form of BOLD correction in SS-SI VASO has been originally been developed, described, and validated for 7T imaging (Huber et al. 2014b) and is based on the assumption that the VASO *T_1_*-contrast (in the *M_z_*-direction) is completely orthogonal to the BOLD 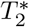-contrast (in the *M_xy_*-direction). In order to minimize noise amplification in the division operation, the BOLD correction was applied on run-averaged time series. Activation GLM analyses were conducted using FSL FEAT (Version 6.0.0). The Feat design used here is available on Github: https://github.com/layerfMRI/repository/tree/master/3T_VASO_scripts/Feat_GLM.

#### 2.4.3. ROI selection

Due to the limited sensitivity of submillimeter VASO fMRI at 3T, we used a combined approach of VASO and BOLD to select the regions of interest (ROIs). First, for an initial large-scale ROI selection, we used the BOLD clusters (FSL-feat, GLM result) after spatial smoothing of 1 mm and binarized them. Then, we dilated them further by 2 voxels. This was done to ensure that the ROIs spanned across the entire cortical depth. Based on these large scale pre-selected ROIs, we ultimately performed layerification in voxels that fulfilled the following criteria:

1. The voxels needed to be located in or at GM. Tissue type segmentation was performed manually based on the inherent *T*_1_-contrast of the mean VASO EPI (underlay in Fig. 4).
2. The voxels had to be located in the Brodman area BA4a. This means that we only focused on the lateral side of the hand knob in the precentral gyrus.
3. The signal was extracted from one entire patch of the cortex that had to be connected (without holes) that had to span across all layers of the cortical depth and had to be at least as wide as the cortical thickness. This criterion was chosen to avoid sampling biases of different detection sensitivity across cortical depth.

**(Fig. 3).**
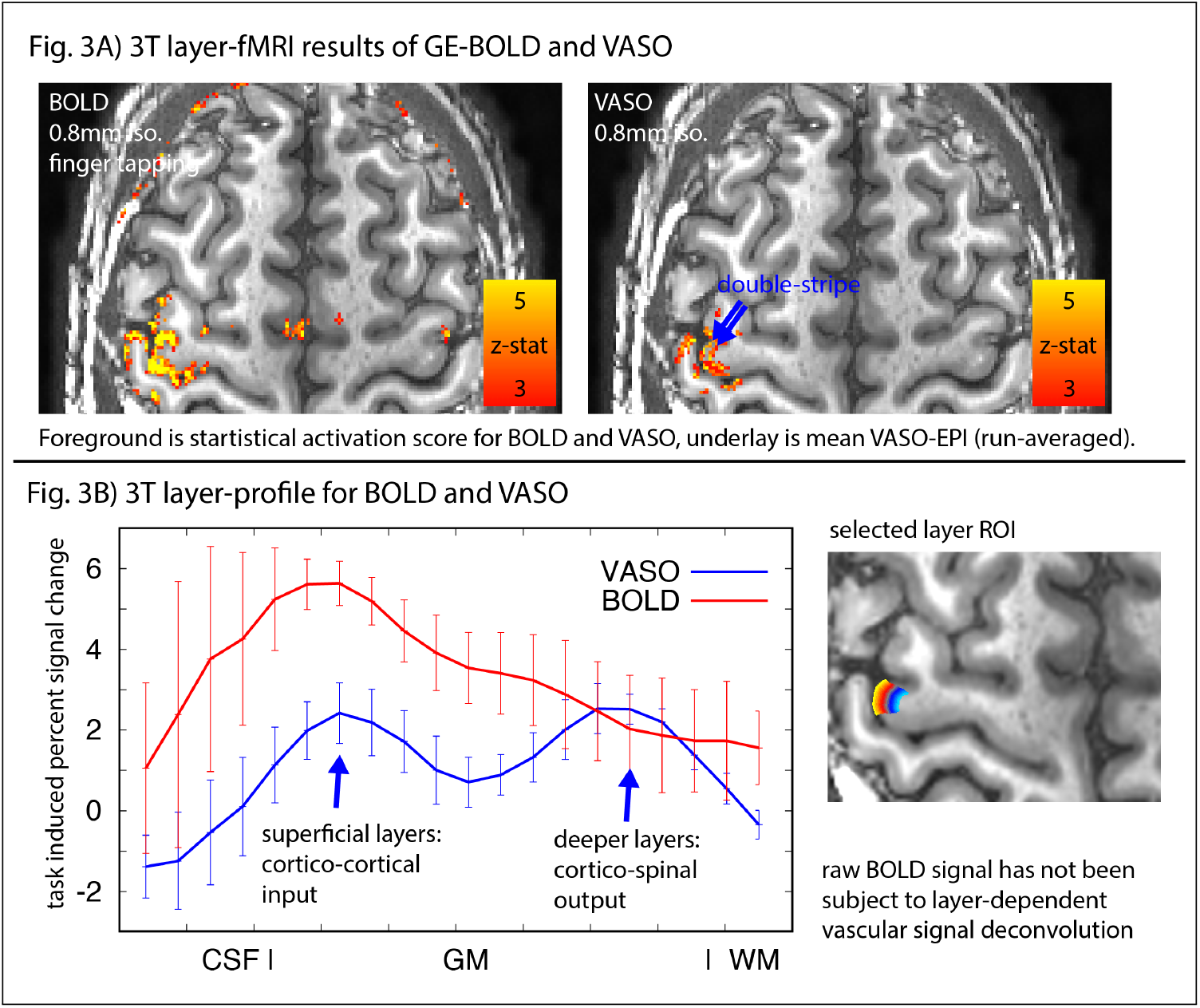
Results of one representative participant. Panel A) depicts the layer-specific detection sensitivity and localization specificity of BOLD and VASO fMRI contrasts at 3T. Indications of a double stripe can be seen (blue arrows). Panel B) depicts the corresponding layer-profiles in the anterior part of the hand knob (BA4A). This approach of brain area-wide voxel averaging has been advocated for BOLD at low field strengths by Markuerkiaga (2020). We believe that this approach is similarly helpful for layer-fMRI VASO. In fact, in current research this approach has enabled layer-fMRI VASO applications in clinical populations (Horovitz et al. 2022).

**Fig. 4).**
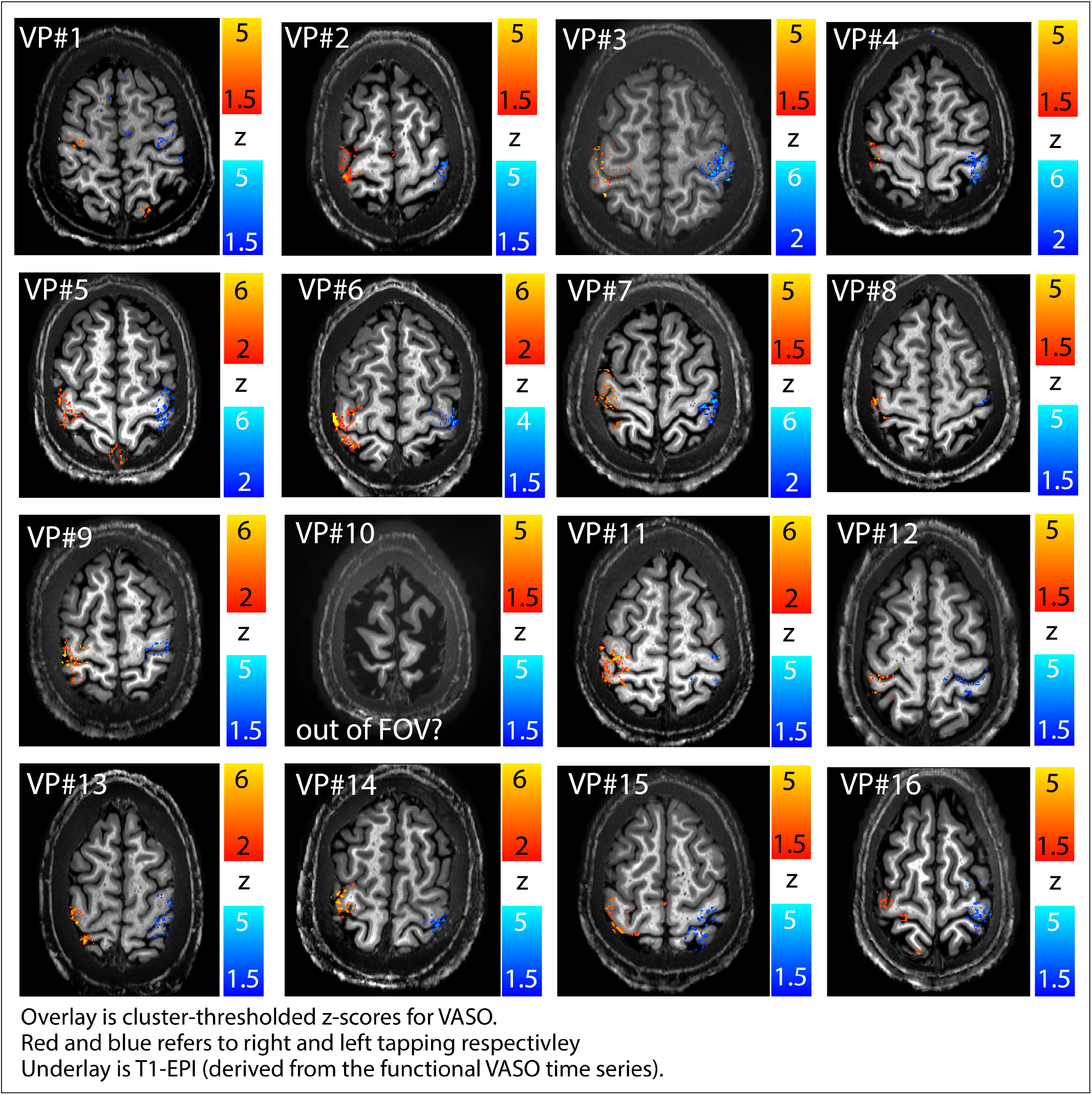
VASO activation maps of all participants (N=16). The data presented here refer to 16 trials acquired across two runs (8 each). Activation patterns can be seen relatively clearly in most participants. We did not find clear activation patterns in one participant (VP10). This is most likely because the imaging slab did not cover the hand area of the primary motor cortex. The data presented here show that the activity pattern mostly follows the GM along the cortical ribbon. The underlay images exemplify the inherent *T*_1_-weighting of the VASO fMRI signal. The panels of this figure also highlight the relatively low artifact level of sub-millimeter, low-bandwidth EPI readout at 3T. Note, that the red and blue colors refer to left-and right-hand finger tapping, which was performed in an interleaved fashion inter-spaced with additional rest periods. Negative CBV responses are not shown in these activation maps. All of the selected 2D slices used in the individual panels are all taken from the inferior part of the slabs, respectivitly.

The exact commands and the order they were executed with are available here: https://github.com/layerfMRI/repository/tree/master/3T_VASO_scripts. Nii files of all ROIs are available for download as part of the shared dataset on the Dataverse repository.

#### 2.4.4. Layerification and layer-signal extraction

Layerfication was performed with the `LN2_LAYERS` program in LayNii (version 2.2.0 (Huber et al. 2021b)) with the equivolume principle (Bok 1929; Waehnert et al. 2014). We included the rim of the GM segmentation as the outermost layers into our analysis. The equivolume depth estimates were binned into equally sized layer groups to contain the same number of voxels across cortical depths. I.e., for a total of 9 layers, the deepest 11% equivolume voxels were assigned to layer 1, the next 11% of the voxels were assigned to layer 2 etc. Note that in the context of fMRI, these estimates of numbered cortical depth do not refer to cytoarchitectonically defined cortical layer numbers. In accordance with the consensus of the field, we still refer to these groups of voxels as ‘layers’ (see https://layerfmri.com/terminology/). This layerification was performed on a spatial grid of 250μm, which is finer than the functional resolution of 0.8mm. This was done to minimize resolution lossed during the binning of voxels into layer groups (https://layerfmri.com/how-many-layers-should-i-reconstruct/).

The functional signal of BOLD and VASO was averaged across all voxels within a given layer group. For interparticipant comparisons the layer profiles were scaled based on their overall response across cortical depth before averaging.

#### 2.4.5. Denoising analysis approaches

To account for the relatively lower expected functional sensitivity at 3T compared to conventional layer-fMRI field strengths, we investigated the effect of post-processing-based denoising by means of layer-specific smoothing. We used the LayNii program `LN_LAYER_SMOOTH` with smoothing kernels between FWHM of 0.4mm and FWHM of 8mm. This smoothing was applied without signal leakage across `–kissing_gyri`. If not explicitly mentioned (e.g. in Fig. 8), no smoothing was applied.

**Fig 5.**
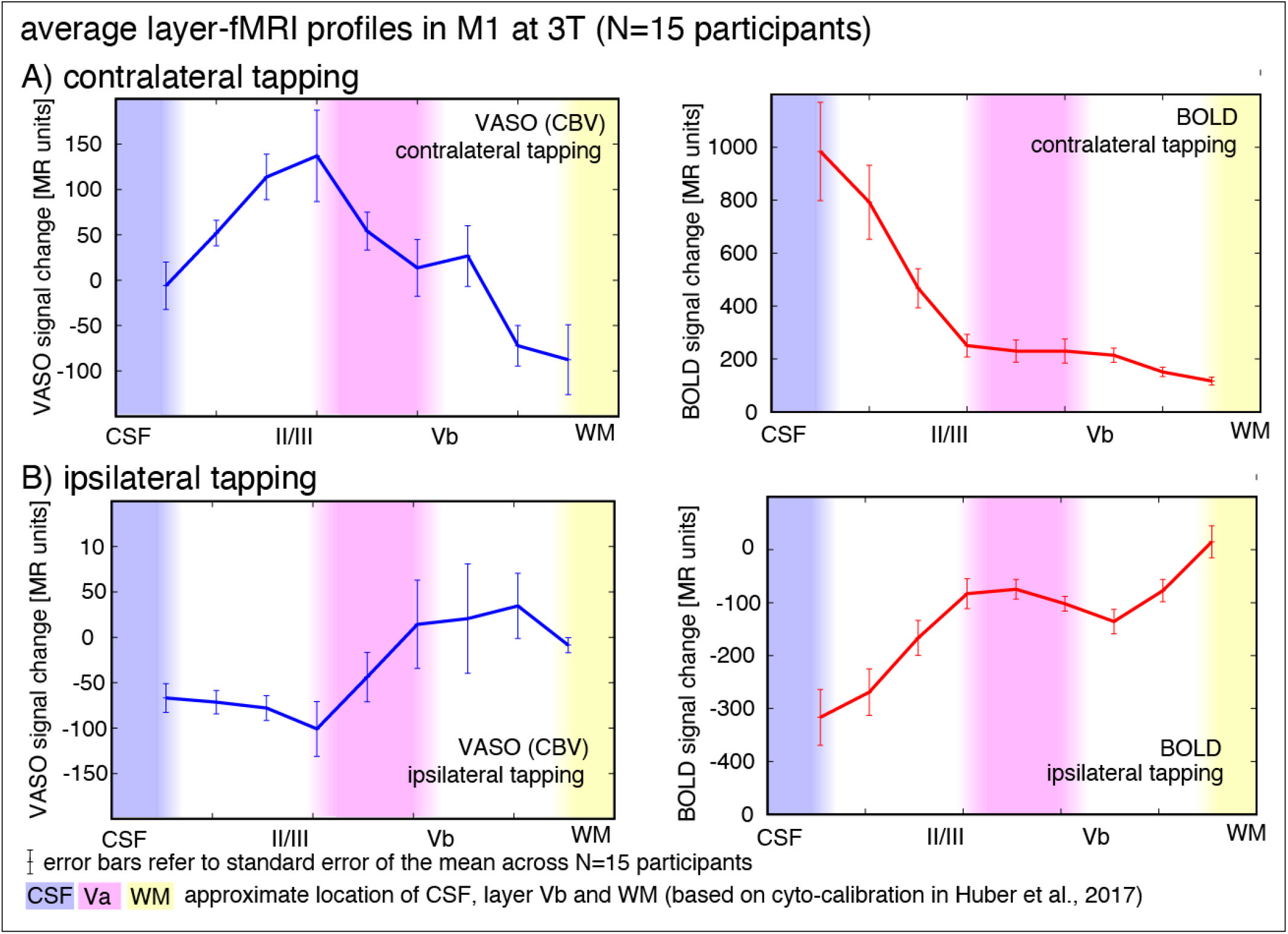
Average layer profiles across participants (N=16). Each data point refers to an average of voxels of a given equivolume cortical depth averaged across participants. The average number of voxels per layer was 81± 24 (Mean± STDEV). Panel A) depicts functional responses during contralateral finger tapping. It can be seen that GE-BOLD is contaminated by inflated signal changes at the surface, whereas VASO profiles peaks in layers II/III.The location of layers is estimated based on depth calibrations using ex-vivo histology (Huber et al. 2017). While the largest CBV responses are located in layers II/III, there are indications of a secondary ‘bump’ (shoulder) in layer Vb/VI. Panel B) depicts functional responses during ipsilateral finger tapping. All profiles are extracted from the same set of voxels, respectively.

**Fig. 6.).**
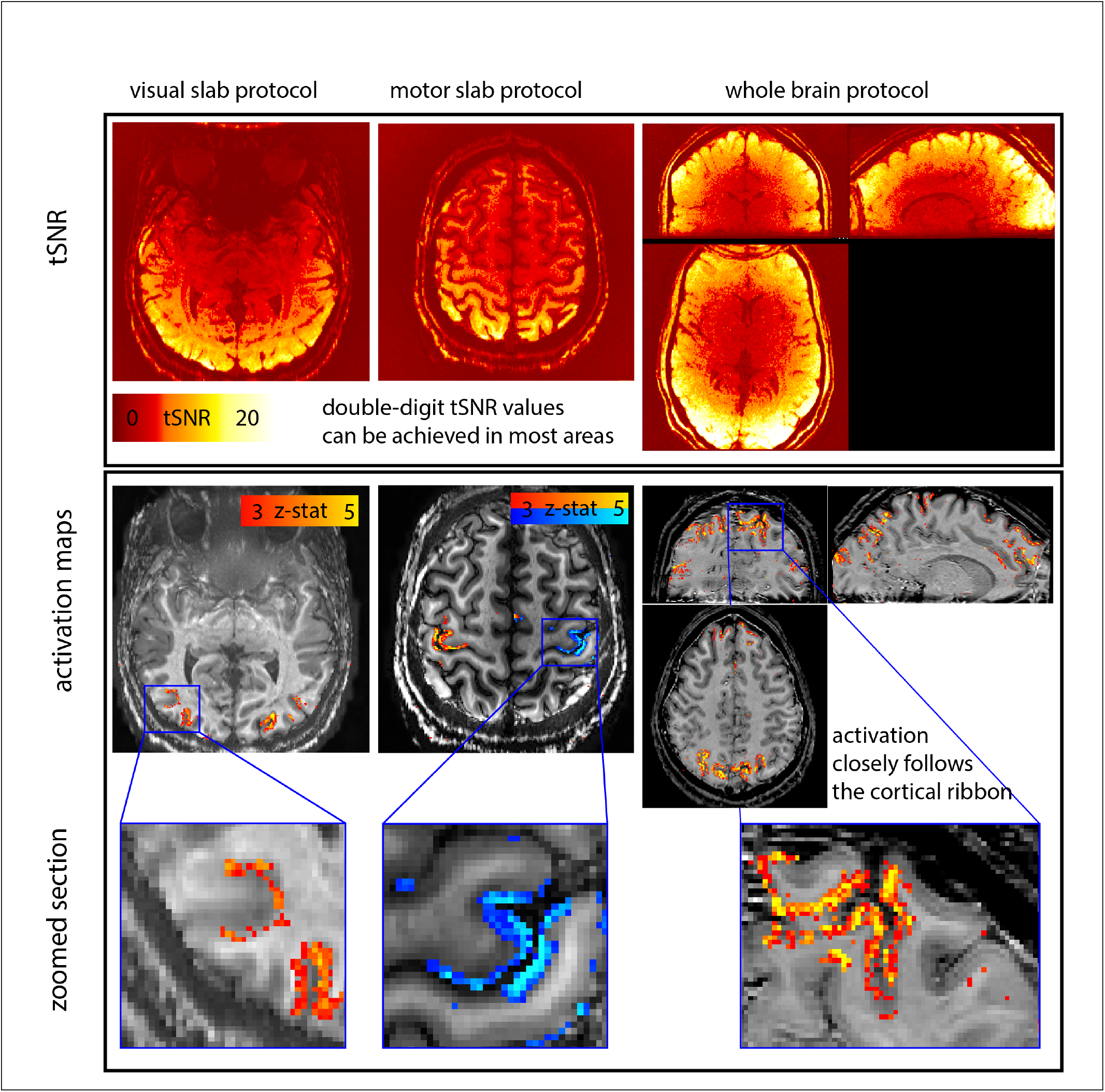
Representative result of exploratory study across protocols. Blood volume weighted fMRI signal quality and activation maps of three layer-fMRI protocols tested. It can be seen that the tested protocols provide enough data quality and signal stability to obtain reasonable activation maps.

**Fig. 7.**
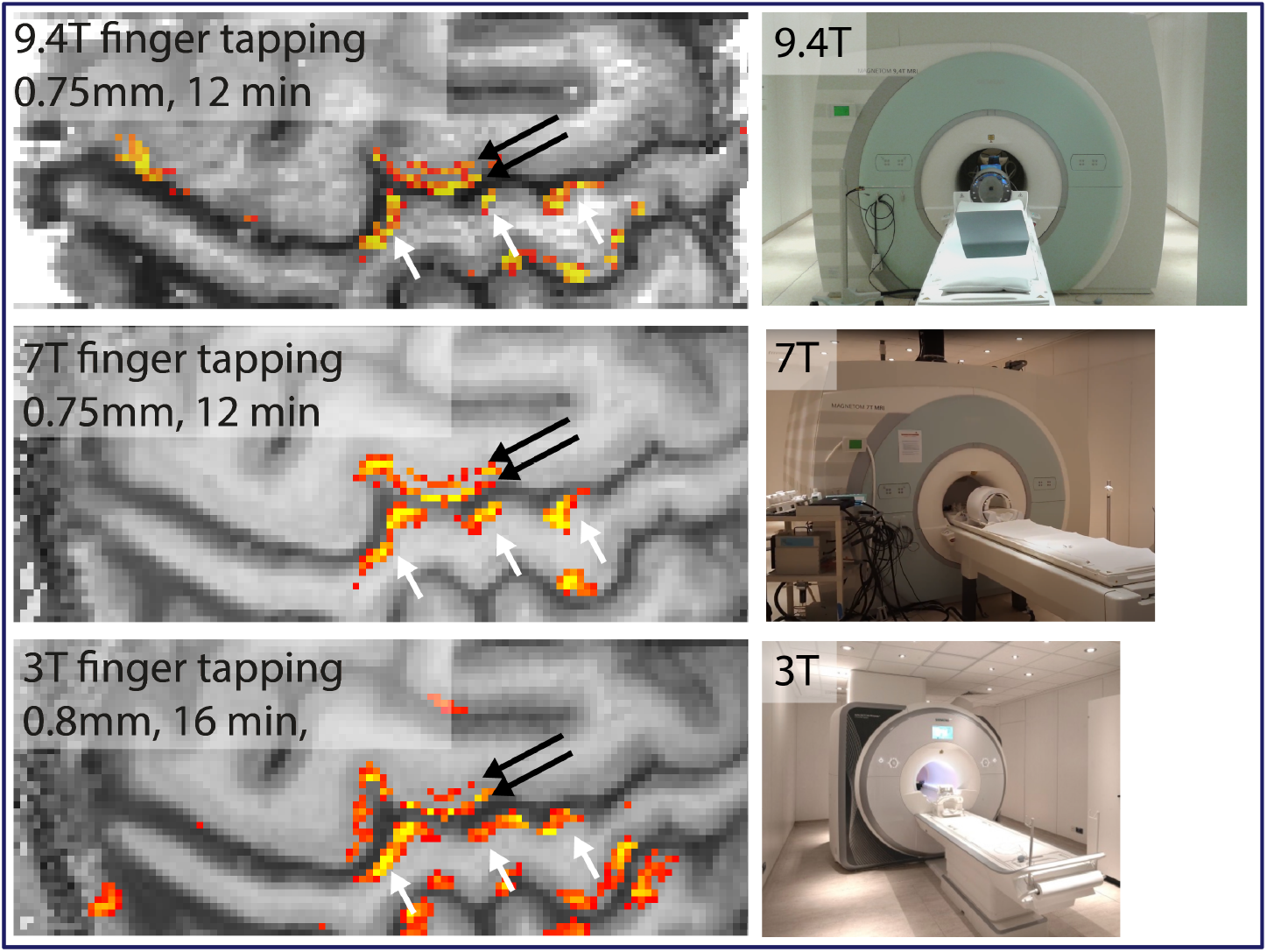
Qualitative comparison across field strength in one participant. Results from the same participant with a very small FOV (32×96 matrix). Despite each field strengths specific challenges, layer-fMRI VASO results can be obtained within 16min experiments.

**Fig. 8.**
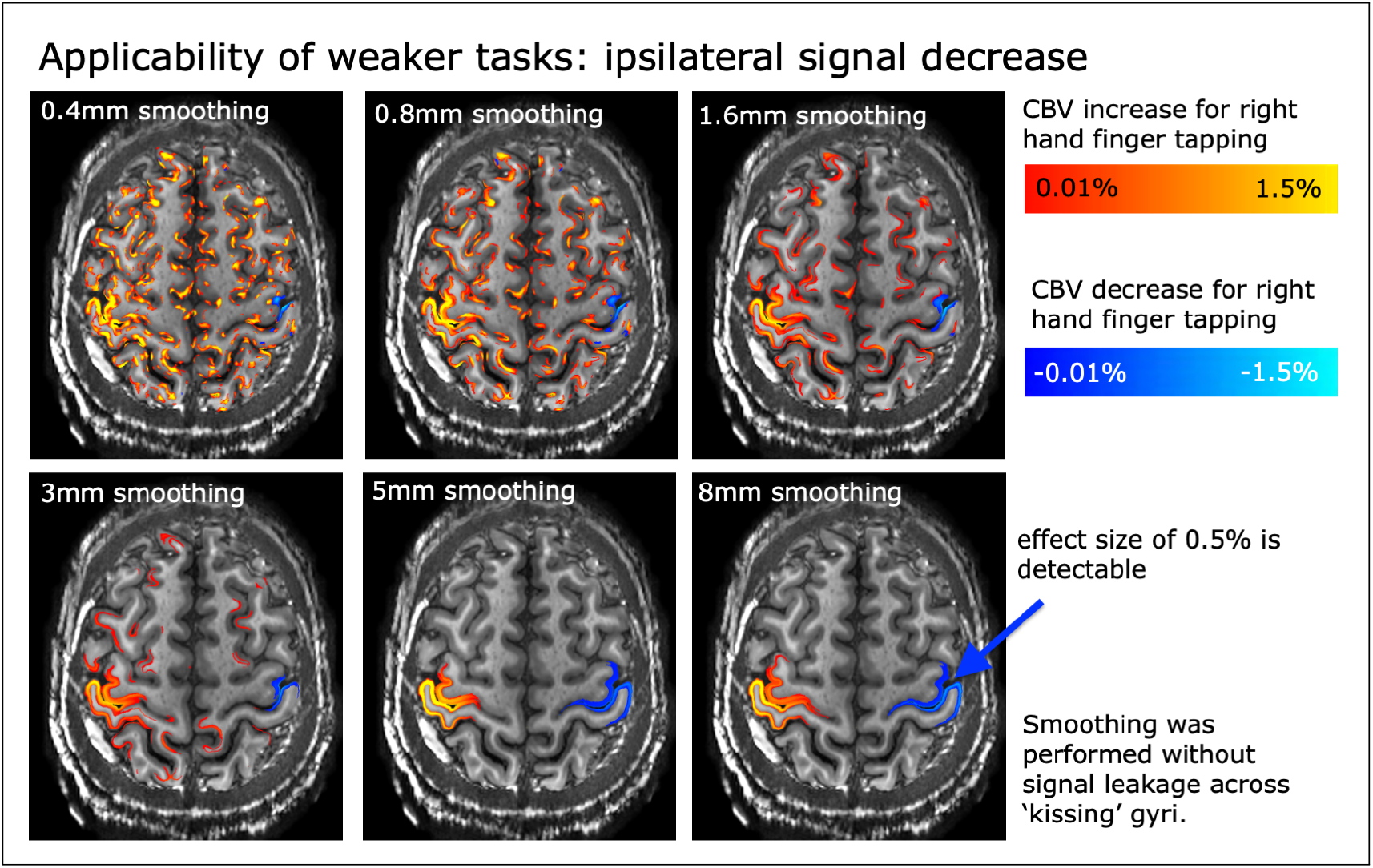
Within-layer smoothing to mitigate the limiting influence of voxel-specific thermal noise. In order to explore the feasibility of layer-fMRI VASO at 3T, we used the weak inhibitory ipsilateral signals as a ‘test-bed’ for any potential weak tasks that are more common for neuroscience application studies. It can be seen that layer-specific smoothing can provide clear activation maps that can capture faint functional signal modulations with local effect sizes of half percent. Such layer-specific smoothing comes along with the loss of ‘columnar’ resolution. Note that the functional contrast shown here solely refers to the tapping induced activity of the right hand. Red and blue colors refer to positive and negative CBV changes. Left-hand tapping results are not shown here.

## 3. Results

Fig. 3A depicts a single run for BOLD and VASO in one representative participant. The GE-BOLD activation map shows more significantly activated voxels than VASO. This is specifically true in pial voxels that are expected to contain large draining veins. In layer proflie plots, two separate peaks are visible in VASO. These results suggest that 3T layer-fMRI VASO imaging methods can provide sufficient signal stability to obtain activation scores on a voxel-by-voxel level to extract layer profiles from small patches of the pinch-movement representation in the primary motor cortex. The relative signal change in the deeper layers is comparable for VASO and BOLD. This suggests that at 3T both imaging modalities can have a similar detection sensitivity for layer-specific micro-vessels. In the superficial layers that also contain draining veins, BOLD signal changes are stronger than VASO signal changes. This is expected due to the unwanted sensitivity of BOLD to large trans-laminar veins.

Fig. 4 and 5 depict the corresponding results across multiple participants. It can be seen that significant VASO signal changes are visible for both task conditions. In one participant the imaging slab did not cover the hand knob. Each task condition refers to 8 trials averaged across two runs. These results suggested that for a single task condition, ≈16 min worth of data are sufficient to obtain reliable sub-millimeter activation maps. The group average layer-profiles in Fig. 5 show that VASO is less sensitive to the surface compared to BOLD. Across participants, the VASO signal changes are strongest in the superficial layers II/III with an indication of a secondary peak in the deeper layers Vb/VI. These two layer groups are expected to be mostly responsible for cortico-cortical input and cortico-spinal output processing, respectively (Mao et al. 2011; Papale and Hooks 2018; Weiler et al. 2008) and they are both expected to be engaged during a tapping task.

Fig. 6 illustrates representative results for the various different tested task protocols: visual protocols, motor protocols and movie watching paradigms. Across all acquisition setups, we found significant activity changes. Due to the local specificity of CBV-fMRI, the activation pattern closely follows the cortical ribbon. This figure also shows tSNR values across brain areas. Most parts of the neocortex have tSNR values in the double-digit regime (>10). The reduced tSNR values in central brain areas are likely due to the larger distance of the small receive elements of the 64ch coil, as well as the larger g-factor penalty with the undersampling patterns used here.

In order to compare the data quality across typical experimental layer-fMRI VASO setups of different field strengths, one participant was asked to repeat the same finger-tapping task at 3T (32ch coil), at 7T (32ch coil), and at 9.4T (31ch coil). Fig. 7 shows thresholded activation maps of these field strength comparisons. These results suggest very similar usability of layer-fMRI across field strengths. It can be seen that the spatial pattern of activated voxels is very similar. E.g., the same double stripe pattern in the hand knob can be seen (black arrows), as well as the inter-space patches in the somatosensory cortex (white arrows).

Fig. 8 shows activation maps with layer-specific smoothing (Blazejewska et al. 2019). It can be seen that this form of denoising, without signal leakage across cortical depth, increases the detection sensitivity so much that it can reveal weak signal responses. In this exploratory study, the weak inhibitory fMRI signal change of 0.5% is used as a representative ‘test-bed’ for any kind of weak effect sizes that are more common in many neuroscience-focused fMRI studies.

Fig. 9 depicts multi-echo results for one representative participant (panels A-B) and participant average results (panel C). These results again support the conclusion of Fig. 3–4, 6–7. Namely, layer-fMRI VASO at 3T provides sufficient signal stability to capture CBV changes in the primary visual and sensorimotor system of the brain. Here, it is shown over a wide range of echo times (12ms, 48ms). It can be seen that the BOLD signal change increases with echo times, whereas VASO signal changes are the same across echo times (within error). This independence of echo time can be used to discuss the potential effect of residual BOLD contamination in 3T VASO.

**Fig. 9.**
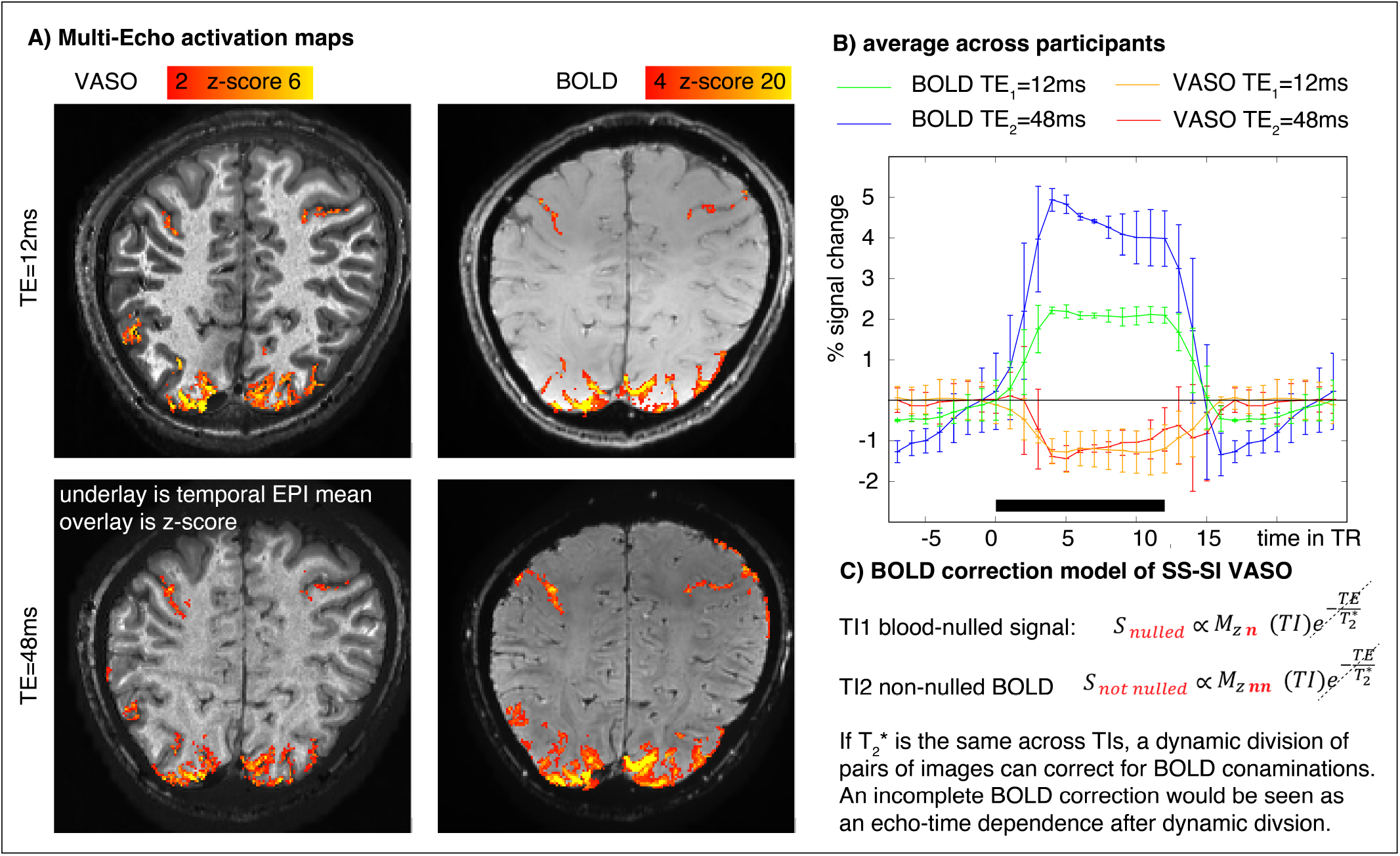
Echo-time dependence of the BOLD and VASO signal. Panel A) depict representative multi-echo activation maps for BOLD and BOLD-corrected VASO. Panel C) shows the participan-averaged time course across echo times. While the BOLD signal change increases with echo times, the VASO signal time course is not different (within error) across echo times. This suggests that for the resolutions, echo times, and voxel selection criteria tested here, the common BOLD correction approach of dynamic division is applicable. Panel D) summarizes the inherent assumption of the BOLD correction method by means of dynamic division. Namely, it is assumed that the modulation of 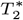 is conserved across inversion times.

## 4. Discussion

The results presented here show that layer-fMRI VASO at 3T is feasible. As expected, we find that the VASO contrast is largely insensitive to unwanted contributions from large draining veins at the cortical surface. This is in line with previous layer-fMRI VASO experiments at 7T (Akbari et al. 2021; Beckett et al. 2020; Persichetti et al. 2020).

### 4.1. Possibility of shorter scan durations or weaker tasks

We show that 16 one-minute runs of finger tapping are sufficient to obtain relatively consistent activation maps with voxel-wise estimations of activation scores. While these results on the feasibility of layer-fMRI at 3T are encouraging, we want to stress that they should be understood as a proof of principle. Future application of layer-fMRI VASO at 3T will likely focus on tasks that come along with smaller effect sizes. Furthermore, future neuroscience-driven studies will likely exploit more task conditions per experiment, which effectively reduces the scan time that can be spent per task condition. In order to make use of 3T layer-fMRI VASO in those more challenging experimental conditions, future studies can employ additional methodological steps to improve on the limited sensitivity of layer-fMRI VASO at 3T.

#### 4.1.1. Layer-specific smoothing

In Fig. 8, we explored the feasibility of layer-specific smoothing in order to capture weak task-evoked VASO signal changes, as they are common in many neuroscience-driven tasks. I.e., we found that faint CBV decreases in the ipsilateral motor cortex during a finger tapping task can be extracted with layer-specific smoothing strengths of ≈ 3mm.

#### 4.1.2. Signal pooling from large ROIs

In many neuroscience-driven applications of layer-fMRI, it is not necessary to resolve activation scores in individual voxels. Instead, some researchers solely aim to investigate brain-area-wide hypotheses. This means that many voxels from a given layer-group can be averaged across large patches of the brain, effectively mitigating thermal noise constraints of layer-fMRI VASO.

#### 4.1.3. Denoising tools

The limited detection sensitivity of layer-fMRI VASO at 3T is expectedly dominated by thermal noise (in contrast to physiological noise and other signal clutter(Wald and Polimeni 2017)). This means that denoising methods that aim to reduce spatio-temporal gaussian noise might be particularly effective here. One very promising denoising candidate is NOise Reduction with DIstribution Corrected (NORDIC) PCA (Vizioli et al. 2021). This approach of NORDIC-PCA denoising for layer-fMRI at 3T has been advocated for BOLD fMRI protocols by Knudsen et al., 2022. First investigations of ongoing research indicated that this approach is similarly helpful for layer-fMRI VASO. In fact, in pilot studies, NORDIC-PCA denoising has allowed 3T layer-fMRI VASO to shorten scan durations by a factor of three, while still maintaining comparable activation maps (Huber et al. 2022).

#### 4.1.4. Advanced hardware

An alternative approach to increase the tSNR is to exploit advanced RF and gradient hardware. For example Akin and Özen showed that microstrip insert head coils allow high-resolution layer-fMRI at 3T within relatively short scan times (Akin and Özen 2019). Furthermore, high-performance head gradients have been proven to be extremely advantageous for layer-fMRI (Feinberg et al. 2022; Polimeni et al. 2010). Thus, we believe that also at lower field strengths, such gradients will be helpful to increase the SNR and, thus, allow shorter scan durations.

#### 4.1.5. Exploring sensitivity of layer-fMRI VASO across field strength

Figure 7 shows tapping induced CBV changes at 3T, 7T, and 9.4T. The purpose of these experiments was to see if there are obvious qualitative differences in the activation maps across field strengths. We find that the spatial pattern of activated voxels are visually similar to the naked eye. The sample size (N=1), the different acquisition duration, the slightly different resolution, the fundamentally different RF coil setup, and the different gradient hardware render quantitative field strengths impossible. However, the results still qualitatively suggest the feasibility of obtaining reproducible activation patterns across field strengths.

### 4.2. Layer-fMRI VASO at 3T enables additional studies that cannot be straightforwardly conducted at higher field strengths

The extension of high-resolution VASO from 7T to more readily available 3T scanners does not only help with the dissemination of laminar imaging methods to a larger user base. More importantly, it allows layer-fMRI applications in experimental setups that have not been possible at 7T before. As such, application studies that are challenged by field inhomogeneity and SAR constraints at 7T, can now be explored at 3T. Examples are:

- **MT-prepared VASO:** Magnetization transfer prepared VASO has been proposed to boost the VASO CBV sensitivity in feeding arteries (Hua et al. 2009; 2013). However at 7T, the conservative SAR restrictions can prohibit a full exploitation of this methodology. We believe that the 3T protocol developed here will be able to mitigate such SAR constraints. At 3T, the layer-fMRI VASO will be able to benefit from such approaches more than at the SAR-restricted UHF field strengths.
- **Brain areas with severe susceptibility artifacts:** At 3T, susceptibility artifacts are significantly reduced compared to higher field settings. This is particularly relevant for brain areas that are inhomogeneity-limited at UHFs. One example is the hippocampus, which is located right above the air-filled ear canals. While there are specific neuroscientific hypotheses with predictions of memory encoding and memory retrieval across the layers in hippocampus, previous layer-fMRI studies of the hippocampus at 7T have been extremely difficult (Chang et al. 2022; Deshpande et al. 2022; Koster et al. 2018; Maass et al. 2014; Zhang et al. 2022). Due to the reduced susceptibility artifacts at 3T, we believe that developments shown in this study will help facilitate more straightforward applications of layer-fMRI in susceptibility-challenged brain areas.
- **Protocols that are limited by physiological noise:** 7T layer-fMRI VASO can be limited by MR-phase inconsistencies that are dynamically modulated during the multi-shot 3D-EPI readout. Such artifacts of layerfMRI VASO have been reported in areas that are particularly challenged by proximal macrovessels and substantial physiological noise like in the primary auditory cortex (Faes et al. 2022). Furthermore, Such artifacts have prohibited the neuroscientific usability of some whole brain layer-fMRI VASO studies with readout durations that are longer than the respiration cycle (Mueller et al. 2021). At 3T, such phase inconsistencies are mitigated. Allowing a more straightforward application of layer-fMRI VASO.
- **Areas with short arterial arrival times:** The SS-SI VASO approach is based on the assumption that all the blood in the imaging slab has been inverted once -and only once. This assumption can be violated for brain areas with very short arterial arrival times, like the insula cortex (Mildner et al. 2014). Furthermore, this assumption can be violated for strong systemic interventions, like *CO*_2_ breathing manipulations (Bright et al. 2011; Chen and Pike 2010). In such cases, fresh, uninverted blood can result in CBF-dependent inflow effects. Without large-scale body transmit RF coils available at 7T due to corresponding SAR constraints, such experimental setups are not straightforwardly usable for VASO. At 3T, however, large transmit coils with lengths of ≥ 50cm are the norm. Thus, we believe that the proposed imaging protocols developed here will facilitate layer-fMRI applications in brain areas like the insula cortex and task modulations like *CO*_2_ breathing.
- **Deep brain structures:** While layer-fMRI has been proven to be valuable for investigating the information flow between cortical brain areas. Many hypotheses about laminar connectivity, however, make predictions about interactions between both, subcortical and cortical regions. In fact, a large portion of cortical input emerges from subcortical brain areas. At 7T, however, the location of subcortical brain areas at the border of the conventional transmit RF coil fields and the distance from the receive channels did not allow cortico-subcortical connectivity studies with layerfMRI VASO. At 3T, the larger body coil for RF transmission and the longer RF wavelength for reception mitigate these constraints. Thus, we believe that the described sequence setup developed here will in the future allow laminar fMRI with connectivity to the subcortex.
- **Multi-modal imaging:** Layer-fMRI can provide complimentary information of neural processing to other neuroimaging modalities including: electroencephalography (EEG), transcranial magnetic stimulation (TMS), or positron emission tomography (PET).

– Concurrent layer-fMRI and EEG can help understanding the neural code of feed-forward and feedback signals (Marsh et al. 2020).
– Concurrent layer-fMRI and TMS has the potential to cross-validate each other regarding differentiating bi-directional or uni-directional connectivity from and to a given node of a brain network (Handwerker et al. 2020).
– Concurrent layer-fMRI and PET, can provide complimentary measures of effective functional connectivity of large brain networks in rest and disease (Riedl et al. 2016).

Such concurrent imaging setups are almost exclusively confined to 3T scanners. The developed protocols here facilitate future multimodal studies with layer-fMRI.
- **Multi-center studies** The field of fMRI is currently experiencing the trend to larger and larger cohort studies. This usually involves participants being acquired as multiple imaging centers. With the limited number of UHF scanners installed, layer-fMRI at 7T cannot follow this trend as easily as conventional low-resolution fMRI. The protocols developed here can pave the road to help future multicenter studies across more readily available 3T scanners.

### 4.3. Value of 3T layer-fMRI VASO besides BOLD

Previous layer-fMRI studies have been conducted at 3T (Akin and Özen 2019; Kim and Ress 2017; Knudsen et al. 2022; Koopmans et al. 2010; Lifshits et al. 2018; Markuerkiaga et al. 2020; Olman et al. 2007; Puckett et al. 2016; Ress et al. 2007; Scheeringa et al. 2016; Taso et al. 2021; Wu et al. 2018). Those studies were somewhat limited by unwanted sensitivities of large draining veins. Here, we furthermore show the feasibility of the non-BOLD contrast, VASO. This contrast can be useful in cases where conventional BOLD signals are hard to interpret due to their spatially unspecific sensitivity to large training veins. In such experiments, utilizing the developed VASO protocols alongside with BOLD can be useful to augment the understanding of the precise depth-dependent location of the fMRI signal origin. Other example studies, where acquiring VASO and GE-BOLD simultaneously may be beneficial, might be related to research questions of altered vascular baseline physiology (Chen 2019) (e.g. in studies about pharmacological interventions, aging, and surgical interventions, etc.). Such experiments are conventionally being conducted at 3T and, thus, have not been investigated on a laminar level yet. The imaging protocol developed here has the potential to facilitate such studies in the future. Furthermore, we think that the concomitantly acquired VASO and BOLD data can be useful to calibrate existing layer-fMRI BOLD models (Corbitt et al. 2018; Havlicek et al. 2015; Heinzle et al. 2016; Markuerkiaga et al. 2016; Merola and Weiskopf 2018). These models have been developed for higher field strengths and are not yet directly applicable for 3T layer-fMRI BOLD data. For example, future GE-BOLD studies that want to apply venous-deconvolution model-inversion at 3T (Puckett et al. 2016) would need to be re-calibrated. The developed imaging protocol and the data it provides have the potential to be informative in this endeavor.

### 4.4. Potential for residual BOLD contamination

With any finite echo time, blood-nulled (VASO) EPI is contaminated with BOLD signal contributions. To account for the dominating extra vascular BOLD effect, layer-fMRI VASO signals are commonly acquired concomitantly with BOLD. This method is known as the SS-SI VASO variant. Measuring the simultaneously ongoing BOLD signal changes, they can be quantified and removed from the VASO signal traces of each voxel. This removal is conducted by means of a dynamic division (Huber et al. 2014b) and can solely account for extra-vascular BOLD. Intra-vascular BOLD cannot be accounted for by this approach and will result in a CBV overestimation. This approach of BOLD correction with dynamic division has been validated by means of investigating the echo time dependency of the BOLD-corrected VASO signal in previous studies at 3T and 7T. Simulation studies assuming a rodent vascular network predicted a residual BOLD contamination of ≈0.2% at 3T and ≈0.15% at 7T, respectively (Genois et al. 2020). Previous empirical multi-echo data in humans, however, found a considerably smaller residual BOLD signal contamination of <0.003% at 7T for TEs between 12 ms and 52 ms (Huber et al. 2014a;b). At 3T, low-resolution (3×3×4mm) multi-echo results found no evidence of any residual echo time dependence in the BOLD corrected VASO signal for echo times between 14ms and 30ms (Huber et al. 2014b). Here, we further investigated the echo-time dependence at 3T with sub-millimeter resolutions across an TE range of 12ms - 48ms (Fig. 8). In significantly activated VASO voxels, we do not find evidence for CBV overestimation with increasing TEs. The lack of CBV overestimation with increasing TEs in this study is somewhat unexpected. At 3T, it is expected that 30% of the overall BOLD signal change is arising from intravascular signal sources (Donahue et al. 2011). Ongoing 3T VASO with multi-echo DEPICTING (Hetzer et al. 2011) suggests that there indeed is a residual BOLD contamination in VASO (Devi et al. 2022). It has been hypothesized that these residual BOLD contaminations might be arising from voxels of large extra-cortical veins. These veins are located outside the GM parenchyma and do not affect the layer-fMRI signal investigated here.

The presence of residual BOLD contaminations in VASO allows future users of this imaging methodology to exploit an additional degree of freedom. Namely, future users can adjust their echo time of choice such that it fits the desired sensitivity-specificity compromise of the functional data. A shorter echo time will have negligible BOLD contaminations and provide functional data that are most locally specific, but less sensitive. A longer echo time will provide an additional sensitivity boost to the SS-SI VASO signal with additional (less specific) BOLD signal contributions.

## Summary and Significance

Examining human cortical layers provides insights into directional information flow of cortical processes, which can otherwise only be achieved via invasive methods like intracortical electrodes. However, while previous layer-fMRI studies show promising findings at ultra-high field strengths, layerfMRI acquisition protocols are commonly confined to specialized research hubs with dedicated ultra-high field centers, thus limiting the useability of laminar fMRI to methods-focused research groups. Extending layer-fMRI VASO to 3T in this study has the potential to make these research tools available to a much wider community. Specifically, the feasibility of sub-millimeter VASO at 3T, allows more neuroscience- and psychiatry-focused research groups to address questions of laminar neural processing in health and disease.

The results presented here taken together suggest that layer-specific CBV-fMRI data can be acquired on widely available clinical 3T scanners. While the functional detection threshold is limited, we show that layer-dependent activation can be visualized in reasonable scan times of 16 min (primary brain areas) to 45 min (associative brain areas). Due to the vast availability of 3T compared to UHF scanners, we believe that this demonstration of 3T-optimized VASO development can boost the user base of laminar fMRI and helps pave the way for CBV layer-fMRI to become a routine tool to address both clinical and basic neuroscience research questions.

## Further material and Data and Software availability

- For a practical “hands-on” perspective of these experiments in the lab, see a 5 min description of the experiments: https://youtu.be/rKjJL-P1mf8. For a 10 min overview video of the experiments conducted here, see: https://youtu.be/cB6v0Dn2FkI.
- Raw and processed data of the main experiment series (N=16) are available here: https://doi.org/10.34894/SRT8RU.
- Data of the multi-echo experiments are available for download here: https://layerfmri.page.link/ME_VASO3T.
- We are happy to share the sequence binaries via SIEMENS’ core competence partnership (C^2^P) agreements.
- An extensive application-focused description of the sequence and how to use it can be found in the form of an FAQ page here: https://layerfmri.com/vaso_ve/.
- Sequence protocols that can be imported to other **SIEMENS** scanners (exar1 files) are available here: https://github.com/layerfMRI/ Sequence_Github/tree/master/3T.
- Layerification analysis code developed in-house is available on Github (https://github.com/layerfMRI/LAYNII).
- Specific application script used for the ROI selection and layerification, are available here: https://github.com/layerfMRI/repository/tree/master/3T_VASO_scripts.

## Acknowledgements

**Advice and inspiration:** We thank Emily Finn for guidance regarding the application of naturalistic tasks. We thank Ratna Devi, Toralf Mildner and Harald E Möller for inspiring us to conduct the multi-echo experiments (Fig. 9) and for multiple discussions on the BOLD correction methods at 3T. We thank Luca Vizioli for advice regarding the application of NORDIC-PCA. We want to thank Paul Taylor for his advice to also discuss the applicability of weaker task effect sizes (inspiring the inclusion of section 3.1 and Fig. 8). We thank Simon Robinson for teaching us details of the specific adaptive coil combination used in IcePat.

## Scanning and sequence

We thank SIEMENS Healthineers for support with sequence code sharing via C2Ps. We thank SIEMENS Healthineer Erik van den Berg for guidance regarding the Prisma transmit coil dimensions. We thank Scannexus for placing their Prisma to our disposal for phantom sequence testing of the protocols used here. We thank the faculty of psychology and neuroscience of the Maastricht University for supporting the multi-echo experiments via scan hours of the ‘development time’.

## Tasks

Movie stimulus data were provided by the Human Connectome Project, WU-Minn Consortium (Principal Investigators: David Van Essen and Kamil Ugurbil; 1U54MH091657) funded by the 16 NIH Institutes and Centers that support the NIH Blueprint for Neuroscience Research; and by the McDonnell Center for Systems Neuroscience at Washington University.

## Diversity Statement

Recent work in several fields of science has identified a bias in citation practices such that papers from women and other minorities are under-cited relative to the number of such papers in the field (Dworkin et al. 2020). In the human layer-fMRI community, the average gender citation bias is 84% male, 15% female (https://layerfmri.com/papers/). For the reference list of this paper, we obtained the gender of the first author of each reference. By this measure (and excluding self-citations to all authors of our current paper), our references contain 47 (70%) male first and 20 (30%) female first. We look forward to future work that could help us to better understand how to support equitable practices in science.

A WHO report from 2008 stated that 90% of the world’s population does not have access to MRI. Since then, the WHO reported a significant increase of scanner density in emerging economies (Geethanath and Krishan 2019). The increasing number of MRI scanners mostly refers to field strengths lower than either 3T or 7T. While the methodology developed here remains to be solely applicable in privileged research centers with 3T scanners, we see this work of porting 7T methods to 3T as a first step into the direction of making these research tools available to a wider more diverse user base.

## References

Akbari, A., Bollmann, S., Ali, T. S., and Barth, M. (2021). Modelling the depth-dependent VASO and BOLD responses in human primary visual cortex. bioRxiv, page 2021.05.07.443052.

Akin, B. and Özen, A. C. (2019). Microstrip Array Insert for Head Coils: Towards Layer fMRI at High Fields. In ISMRM, page 0371.

Beckett, A. J., Dadakova, T., Townsend, J., Huber, L., Park, S., and Feinberg, D. A. (2020). Comparison of BOLD and CBV using 3D EPI and 3D GRASE for cortical layer functional MRI at 7 T. Magnetic Resonance in Medicine, 84(6):3128–3145.

Blazejewska, A. I., Fischl, B., Wald, L. L., and Polimeni, J. R. (2019). Intracortical smoothing of small-voxel fMRI data can provide increased detection power without spatial resolution losses compared to conventional large-voxel fMRI data. NeuroImage, 189(January):601–614.

Bok, S. (1929). Der Einfluß der in den Furchen und Windungen auftretenden Krümmungen der Großhirnrinde auf die Rindenarchitektur. Gesamte Neurol. Psychiatr., 12:682–750.

Bollmann, S. and Barth, M. (2020). New acquisition techniques and their prospects forthe achievable resolution of fMRI. Progress in Neurobiology, 207(October):101936.

Breuer, F. A., Blaimer, M., Mueller, M. F., Seiberlich, N., Heidemann, R. M., Griswold, M. A., and Jakob, P. M. (2006). Controlled aliasing in volumetric parallel imaging (2D CAIPIRINHA). Magnetic Resonance in Medicine, 55(3):549–556.

Bright, M. G., Donahue, M. J., Duyn, J. H., Jezzard, P., and Bulte, D. P. (2011). The effect of basal vasodilation on hypercapnic and hypocapnic reactivity measured using magnetic resonance imaging. Journal of Cerebral Blood Flow and Metabolism, 31(2):426–438.

Chang, W.-t., Langella, S., and Giovanello, K. (2022). Cross-layer Balance of Visuo-hippocampal Functional Connectivity Is Associated With Episodic Memory Recognition Accuracy. Research Square, pages 1–21.

Chen, J. J. (2019). Functional MRI of brain physiology in aging and neurodegenerative diseases. NeuroImage, 187(May 2018):209–225.

Chen, J. J. and Pike, G. B. (2010). MRI measurement of the BOLD-specific flow-volume relationship during hypercapnia and hypocapnia in humans. NeuroImage, 53(2):383–391.

Corbitt, P. T., Ulloa, A., and Horwitz, B. (2018). Simulating laminar neuroimaging data for a visual delayed match-to-sample task. NeuroImage, 173(February):199–222.

Deshpande, G., Zhao, X., and Robinson, J. (2022). Functional parcellation of the hippocampus based on its layer-specific connectivity with default mode and dorsal attention networks. NeuroImage, 254(March):119078.

Devi, R., Mildner, T., Schlumm, T., and Möller, H. E. (2022). CBV-Based fMRI at 3T with SS-SI-VASO: Multi-Echo DEPICTING vs Multi-Echo EPI. In Ismrm, page 2111.

Donahue, M. J., Hoogduin, H., Van Zijl, P. C., Jezzard, P., Luijten, P. R., and Hendrikse, J. (2011). Blood oxygenation level-dependent (BOLD) total and extravascular signal changes and ΔR2* in human visual cortex at 1.5, 3.0 and 7.0 T. NMR in Biomedicine, 24(1):25–34.

Dworkin, J. D., Linn, K. A., Teich, E. G., Zurn, P., Shinohara, R. T., and Bassett, D. S. (2020). The extent and drivers of gender imbalance in neuroscience reference lists. Nature Neuroscience, 23(8):918–926.

Faes, L. K., Gulban, O. F., Poser, B. A., Martino, F. D., and Huber, R. (2022). CBV-sensitive layer-fMRI in the human auditory cortex at 7T: Challenges and capabilities. In Ismrm, page 2194.

Feinberg, D. A., Torrisi, S., Beckett, A. J., Stirnberg, R., Stöcker, T., Ehses, P., and Huber, R. (2022). Sub-0.1 microliter CBV fMRI on the Next Generation 7T. In ISMRM.

Geethanath, S. and Krishan, S. (2019). Accessible MRI. Magn Reson Med, 56(1).

Genois, E., Gagnon, L., and Desjardins, M. (2020). Modeling of vascular space occupancy and BOLD functional MRI from first principles using real microvascular angiograms. Magnetic Resonance in Medicine, 85(December 2019):456–468.

Griswold, M. A., Jakob, P. M., Heidemann, R. M., Nittka, M., Jellus, V., Wang, J., Kiefer, B., and Haase, A. (2002). Generalized Autocalibrating Partially Parallel Acquisitions (GRAPPA). Magnetic Resonance in Medicine, 47(6):1202–1210.

Handwerker, D. A., Ianni, G., Gutierrez, B., Roopchansingh, V., Gonzalez-Castillo, J., Chen, G., Bandettini, P. A., Ungerleider, L. G., and Pitcher, D. (2020). Theta-burst TMS to the posterior superior temporal sulcus decreases resting-state fMRI connectivity across the face processing network. Network Neuroscience, 4(3):746–760.

Havlicek, M., Roebroeck, A., Friston, K., Gardumi, A., Ivanov, D., and Uludag, K. (2015). Physiologically informed dynamic causal modeling of fMRI data. NeuroImage, 122:355–372.

Heinzle, J., Koopmans, P. J., den Ouden, H. E., Raman, S., and Stephan, K. E. (2016). A hemodynamic model for layered BOLD signals. NeuroImage, 125:556–570.

Hetzer, S., Mildner, T., and Möller, H. E. (2011). A modified EPI sequence for high-resolution imaging at ultra-short echo time. Magnetic resonance in medicine: official journal of the Society of Magnetic Resonance in Medicine / Society of Magnetic Resonance in Medicine, 65(1):165–175.

Horovitz, S. G., Kassavetis, P., Hallett, M., and Huber, R. (2022). Laminar VASO fMRI in hand dystonia patients. In ISMRM.

Hua, J., Donahue, M. J., Zhao, J. M., Grgac, K., Huang, A. J., Zhou, J., and Van Zijl, P. C. (2009). Magnetization transfer enhanced Vascular-space-occupancy (MT-VASO) functional MRI. Magnetic Resonance in Medicine, 61(4):944–951.

Hua, J., Jones, C. K., Qin, Q., and Van Zijl, P. C. (2013). Implementation of vascular-space-occupancy MRI at 7T. Magnetic Resonance in Medicine, 69(4):1003–1013.

Huber, L., Finn, E. S., Chai, Y., Goebel, R., Stirnberg, R., Stöcker, T., Marrett, S., Uludag, K., Kim, S. G., Han, S. H., Bandettini, P. A., and Poser, B. A. (2021a). Layer-dependent functional connectivity methods. Progress in Neurobiology, 207.

Huber, L., Goense, J., Kennerley, A. J., Ivanov, D., Krieger, S. N., Lepsien, J., Trampel, R., Turner, R., and Möller, H. E. (2014a). Investigation of the neurovascular coupling in positive and negative BOLD responses in human brain at 7T. NeuroImage, 97:349–362.

Huber, L., Handwerker, D. A., Jangraw, D. C., Chen, G., Hall, A., Stüber, C., Gonzalez-Castillo, J., Ivanov, D., Marrett, S., Guidi, M., Goense, J., Poser, B. A., and Bandettini, P. A. (2017). High-Resolution CBV-fMRI Allows Mapping of Laminar Activity and Connectivity of Cortical Input and Output in Human M1. Neuron, 96(6):1253–1263.

Huber, L., Ivanov, D., Krieger, S. N., Streicher, M. N., Mildner, T., Poser, B. A., Möller, H. E., and Turner, R. (2014b). Slab-selective, BOLD-corrected VASO at 7 tesla provides measures of cerebral blood volume reactivity with high signal-to-noise ratio. Magnetic Resonance in Medicine, 72(1):137–148.

Huber, L., Tse, D. H., Wiggins, C. J., Uludağ, K., Kashyap, S., Jangraw, D. C., Bandettini, P. A., Poser, B. A., and Ivanov, D. (2018). Ultra-high resolution blood volume fMRI and BOLD fMRI in humans at 9.4 T: Capabilities and challenges. NeuroImage, 178:769–779.

Huber, L. R. R., Poser, B. A., Bandettini, P. A., Arora, K., Wagstyl, K., Cho, S., Goense, J., Nothnagel, N., Morgan, A. T., van den Hurk, J., Müller, A. K., Reynolds, R. C., Glen, D. R., Goebel, R., and Gulban, O. F. (2021b). LayNii: A software suite for layer-fMRI. NeuroImage, 237.

Huber, R., Kronbichler, L., Stirnberg, R., Vizioli, L., Ehses, P., Stöcker, T., Fernández-Cabello, S., Poser, B. A., and Kronbichler, M. (2022). Evaluating the capabilities and challenges of layer-fMRI VASO at 3T. In ISMRM, page 0398.

Jellúš, V. and Kannengießer, S. A. R. (2014). Improved Coil Combination using the Prescan Normalize Adjustment. In SIEMENS Idea meeting 2014, page 4406.

Kellman, P. and McVeigh, E. R. (2005). Image reconstruction in SNR units: A general method for SNR measurement. Magnetic Resonance in Medicine, 54(6):1439–1447.

Kim, J. H. and Ress, D. (2017). Reliability of the depth-dependent high-resolution BOLD hemodynamic response in human visual cortex and vicinity. Magnetic Resonance Imaging, 39:53–63.

Knudsen, L., Bailey, C. J., Blicher, J. U., Yang, Y., Zhang, P., and Torben, E. (2022). Feasibility of 3T layer-dependent fMRI with GE-BOLD using NORDIC and phase regression. bioRxiv, pages 1–26.

Koopmans, P. J., Barth, M., and Norris, D. G. (2010). Layer-specific BOLD activation in human V1. Human Brain Mapping, 31(9):1297–1304.

Koster, R., Chadwick, M. J., Chen, Y., Berron, D., Banino, A., Düzel, E., Hassabis, D., and Kumaran, D. (2018). Big-Loop Recurrence within the Hippocampal System Supports Integration of Information across Episodes. Neuron, 99(6):1342–1354.

Lifshits, S., Tomer, O., Shamir, I., Barazany, D., Tsarfaty, G., Rosset, S., and Assaf, Y. (2018). Resolution considerations in imaging of the cortical layers. NeuroImage, 164(February):112–120.

Lu, H., Golay, X., Pekar, J. J., and Van Zijl, P. C. M. (2003). Functional magnetic resonance imaging based on changes in vascular space occupancy. Magnetic Resonance in Medicine, 50(2):263–274.

Maass, A., Schütze, H., Speck, O., Yonelinas, A., Tempelmann, C., Heinze, H. J., Berron, D., Cardenas-Blanco, A., Brodersen, K. H., Stephan, K. E., and Düzel, E. (2014). Laminar activity in the hippocampus and entorhinal cortex related to novelty and episodic encoding. Nature Communications, 5.

Mao, T., Kusefoglu, D., Hooks, B. M., Huber, D., Petreanu, L., and Svoboda, K. (2011). Long-Range Neuronal Circuits Underlying the Interaction between Sensory and Motor Cortex. Neuron, 72(1):111–123.

Markuerkiaga, I., Bains, L. J., Marques, J. P., Norris, D. G., Bains, L. J., and Norris, D. G. (2020). An in-vivo study of BOLD laminar responses as a function of echo time and static magnetic field strength. Scientific Reports, 11(0123456789):1–14.

Markuerkiaga, I., Barth, M., and Norris, D. G. (2016). A cortical vascular model for examining the specificity of the laminar BOLD signal. NeuroImage, 132:491–498.

Marsh, D., Sokoliuk, R., and Mullinger, K. (2020). Assessing the origin of human alpha oscillations using laminar layer 7T fMRI-EEG. In ISMRM, page 1345.

Martindale, J., Kennerley, A. J., Johnston, D., Zheng, Y., and Mayhew, J. E. (2008). Theory and generalization of Monte Carlo models of the BOLD signal source. Magnetic Resonance in Medicine, 59(3):607–618.

Merola, A. and Weiskopf, N. (2018). Modelling the laminar GE-BOLD signal: integrating anatomical, physiological and methodological factors. In Proceedings International Society for Magnetic Resonance in Medicine, page 2299.

Mildner, T., Müller, K., Hetzer, S., Trampel, R., Driesel, W., and Möller, H. E. (2014). Mapping of arterial transit time by intravascular signal selection. NMR Biomed, 27(February):594–609.

Mueller, A. K., Miriam Heynckes, Wiggins, C. J., Gulban, O. F., Chai, Y., Poser, B., and Renzo Huber (2021). Whole brain layer-fMRI: An open dataset for methods benchmarking. In Ismrm.

Olman, C. A., Inati, S., and Heeger, D. J. (2007). The effect of large veins on spatial localization with GE BOLD at 3 T: Displacement, not blurring. NeuroImage, 34(3):1126–1135.

Papale, A. E. and Hooks, B. M. (2018). Circuit changes in motor cortex during motor skill learning.

Persichetti, A. S., Avery, J. A., Huber, L., Merriam, E. P., and Martin, A. (2020). Layer-Specific Contributions to Imagined and Executed Hand Movements in Human Primary Motor Cortex. Current Biology, 30(9):1721–1725.

Pohmann, R., Speck, O., and Scheffler, K. (2016). Signal-to-noise ratio and MR tissue parameters in human brain imaging at 3, 7, and 9.4 tesla using current receive coil arrays. Magnetic Resonance in Medicine, 75(2):801–809.

Polimeni, J. R., Fischl, B., Greve, D. N., and Wald, L. L. (2010). Laminar analysis of 7 T BOLD using an imposed spatial activation pattern in human V1. NeuroImage, 52(4):1334–1346.

Poser, B. A., Koopmans, P. J., Witzel, T., Wald, L. L., and Barth, M. (2010). Three dimensional echo-planar imaging at 7 Tesla. NeuroImage, 51(1):261–266.

Puckett, A. M., Aquino, K. M., Robinson, P. A., Breakspear, M., and Schira, M. M. (2016). The spatiotemporal hemodynamic response function for depth-dependent functional imaging of human cortex. NeuroImage, 139:240–248.

Ress, D., Glover, G. H., Liu, J., and Wandell, B. (2007). Laminar profiles of functional activity in the human brain. NeuroImage, 34(1):74–84.

Riedl, V., Utz, L., Castrillón, G., Grimmer, T., Rauschecker, J. P., Ploner, M., Friston, K. J., Drzezga, A., and Sorg, C. (2016). Metabolic connectivity mapping reveals effective connectivity in the resting human brain. Proceedings of the National Academy of Sciences of the United States of America, 113(2):428–433.

Scheeringa, R., van Mourik, T., Jensen, O., Koopmans, P. J., and Norris, D. G. (2016). The relationship between oscillatory EEG activity and the laminar-specific BOLD signal. Proceedings of the National Academy of Sciences, 113(24):6761–6766.

Schmitt, T. and Rieger, J. W. (2021). Recommendations of Choice of Head Coil and Prescan Normalize Filter Depend on Region of Interest and Task. Frontiers in Neuroscience, 15(October):1–16.

Stirnberg, R. and Stöcker, T. (2021). Segmented K-Space Blipped-Controlled Aliasing in Parallel Imaging (Skipped-CAIPI) for High Spatiotemporal Resolution Echo Planar Imaging. Magnetic Resonance in Medicine, 85(0):1540–1551.

Taso, M., Munsch, F., Zhao, L., and Alsop, D. C. (2021). Regional and depth-dependence of cortical blood-flow assessed with high-resolution Arterial Spin Labeling (ASL). Journal of Cerebral Blood Flow and Metabolism.

Vizioli, L., Moeller, S., Dowdle, L., Akçakaya, M., De Martino, F., Yacoub, E., and Uğurbil, K. (2021). Lowering the thermal noise barrier in functional brain mapping with magnetic resonance imaging. Nature Communications, 12(1).

Waehnert, M. D., Dinse, J., Weiss, M., Streicher, M. N., Waehnert, P., Geyer, S., Turner, R., and Bazin, P. L. (2014). Anatomically motivated modeling of cortical laminae. NeuroImage, 93:210–220.

Wald, L. L. and Polimeni, J. R. (2017). Impacting the effect of fMRI noise through hardware and acquisition choices – Implications for controlling false positive rates. NeuroImage, 154(December 2016):15–22.

Weiler, N., Wood, L., Yu, J., Solla, S. A., and Shepherd, G. M. (2008). Top-down laminar organization of the excitatory network in motor cortex. Nature Neuroscience, 11(3):360–366.

Weldon, K. B. and Olman, C. A. (2020). Forging a path to mesoscopic imaging success with ultra-high field functional magnetic resonance imaging. Philosophical Transactions B.

Wu, P. Y., Chu, Y. H., Lin, J. F. L., Kuo, W. J., and Lin, F. H. (2018). Feature-dependent intrinsic functional connectivity across cortical depths in the human auditory cortex. Scientific Reports, 8(1):1–14.

Yang, J., Huber, L., Yu, Y., and Bandettini, P. A. (2021). Neuroscience and Biobehavioral Reviews Linking cortical circuit models to human cognition with laminar fMRI. Neuroscience and Biobehavioral Reviews, 128:467–478.

Zhang, K., Chen, L., Li, Y., Paez, A. G., Miao, X., Cao, D., Gu, C., Bakker, A., and Hua, J. (2022). Differential laminar activation in the medial and lateral entorhinal cortex in the human brain revealed by laminar functional MRI at 7T. In ISMRM, High field Workshop.

